# Ciliary protein CEP290 regulates focal adhesion via microtubule system in non-ciliated cells^1^

**DOI:** 10.1101/2023.04.02.535304

**Authors:** Kazuhiko Matsuo, Yoshiro Nakajima, Masaki Shigeta, Daisuke Kobayashi, Shinichiro Sakaki, Satoshi Inoue, Naoki Takeshita, Atsuko Ueyama, Kousuke Nishikawa, Rie Saba, Takahiko Yokoyama, Kenta Yashiro

## Abstract

Almost all differentiated mammalian cells have primary cilia on their surface. Ciliary dysfunction causes ciliopathy in humans. Centrosomal protein 290 (CEP290) is a ciliary protein that causes ciliopathies, localizes at the cilial base in ciliated cells, whereas it localizes to the centrosome in non-ciliated proliferating cells. The cilia-dependent function of CEP290 has been extensively studied; however, the cilia-independent function, which is likely responsible for the wider phenotypic spectra of CEP290-related ciliopathies, remains largely unknown. Here, we examined cilia-independent functions of CEP290 in non-ciliated cells. Our study showed that *Cep290* function loss suppresses microtubule elongation due to microtubule organizing center malfunction. Surprisingly, CEP290 forms a complex with the adenomatous polyposis coli (APC) protein encoded by the *adenomatous polyposis coli* gene. The APC-CEP290 complex exists in the centrosome and on microtubule fibers. Notably, the reduced focal adhesion formation is likely responsible for the *Cep290* mutant phenotypes, including impaired directed cell migration, shrunken cell shape, and reduced adhesive capacity to the extracellular matrix. The APC-CEP290 complex is consistently important for transporting a focal adhesion molecule, paxillin, to focal adhesions in non-ciliated cells. Thus, our findings provide a novel platform to better understand the ciliopathies.

## 1. Introduction

The centrosome is a non-membranous organelle that functions as a microtubule-organizing center (MTOC) and regulates cell polarity, intracellular trafficking, and migration via microtubules in interphase cells. During mitosis, the centrosome organizes the microtubules of the mitotic spindle for faithful chromosome segregation [1]. In differentiated cells and/or cells in the G0 phase, the centrosome moves near the plasma membrane and functions as a basal body that serves as the base for primary cilium formation. Primary cilia are vital for sensing and transmitting extracellular cues to cells to regulate diverse cellular processes. Functional abnormalities of primary cilia cause “ciliopathies” , a group of rare inherited human diseases that usually affect multiple organs [2].

Centrosomal protein 290 (CEP290), a primary cilia protein, localizes to the transition zone at the base of primary cilia [3]. Mutations in this gene are responsible for ciliopathies, including nephronophthisis (NPHP) [4], Bardet–Biedl syndrome (BBS) [5], Joubert syndrome (JS) [3,6], Senior-Løken syndrome (SLS) [7], and Meckel syndrome (MKS) [8,9]. CEP290 is essential for initiating assembly of ciliary transition zone [10], molecular integrity of primary cilia [11], and formation of aggresome [12]. As is generally observed in ciliopathies, the CEP290 phenotype is pleiotropic, such as that observed in polycystic kidney disease, retinitis pigmentosa, and skeletal dysplasia. Although the molecular function of CEP290 has been extensively studied, our understanding of the mechanisms underlying this phenotypic pleiotropy remains incomplete.

CEP290 localizes to the centrosome in non-ciliated cells [6,13,14]. Therefore, CEP290 likely has a cilia-independent function. Here, we hypothesized that the phenotypes caused by CEP290 malfunction originate not only from the primary cilia malfunction but also from microtubule systems dysregulated by the centrosome. The synergistic effect of primary ciliary dysfunction and the microtubule system may be responsible for a wide range of ciliopathic phenotypes. To validate this hypothesis, we attempted to distinguish between these two effects– attributable and not attributable to primary cilia–by focusing on the molecular function of CEP290 in non-ciliated cells. We found that CEP290 localizes not only at the centrosome as previously reported but also on microtubule fibers.

Centrosome/microtubule CEP290 regulates microtubule growth, which is essential for directed cell migration and adhesion. Surprisingly, the adenomatous polyposis coli (APC)-CEP290 complex transports paxillin, a core component of focal adhesions, to the focal adhesion plaques. Our findings provide a novel approach to better understand the pathophysiology of ciliopathies.

## 2. Materials and Methods

### 2.1 The establishment of *Cep290* knockout and PA tagged *Cep290* cell lines

Murine inner medullary collecting duct 3 (mIMCD3) cells (ATCC) were cultured in Dulbecco’s Modified Eagle’s Medium (DMEM)/Ham’s F12 (1:1) medium containing 10% of fetal bovine serum, 100 U/mL of penicillin G and 100 µg/mL of streptomycin.

Two single guide RNA (sgRNAs) were designed to establish the *Cep290* null mutation cell lines using the CRISPR/Cas9 system [15]. The following oligonucleotides were synthesized:

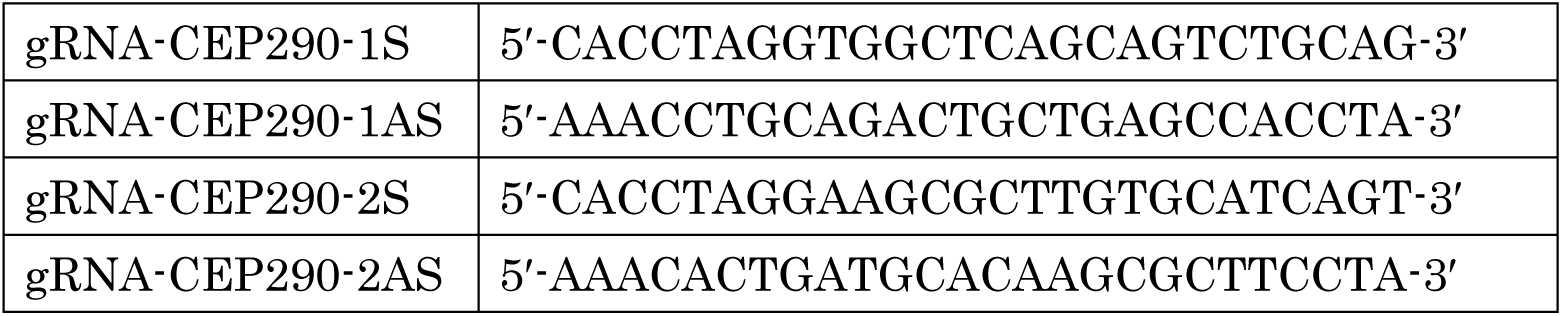

The oligonucleotides of the complementary S and AS strands were annealed, and the 5′-end was phosphorylated with T4 polynucleotide kinase (Cat. # 2021S: Takara, Kyoto, Japan). This phosphorylated DNA fragment was ligated to the *Bbs* I cloning site in pSpCas9(BB)-2A-Puro (PX459) V2.0 (Addgene plasmid # 62988) [16], and the constructs corresponding to 1S/1AS and 2S/2AS were designated as pX459-sgCEP290-KO1 and pX459-sgCEP290-KO2, respectively. pX459-sgCEP290-KO1 or pX459-sgCEP290-KO2 was independently transfected into mIMCD3 cells using Lipofectamine 2000 (Thermo Fisher Scientific, Waltham, MA, USA) according to the manufacturer’s protocol. Puromycin-resistant clones were isolated using 2 μg/mL of puromycin 48 hours after transfection. The genotype of the isolated clone was confirmed by sequencing PCR products from the targeted genomic region. The following primers were used for PCR and sequencing:

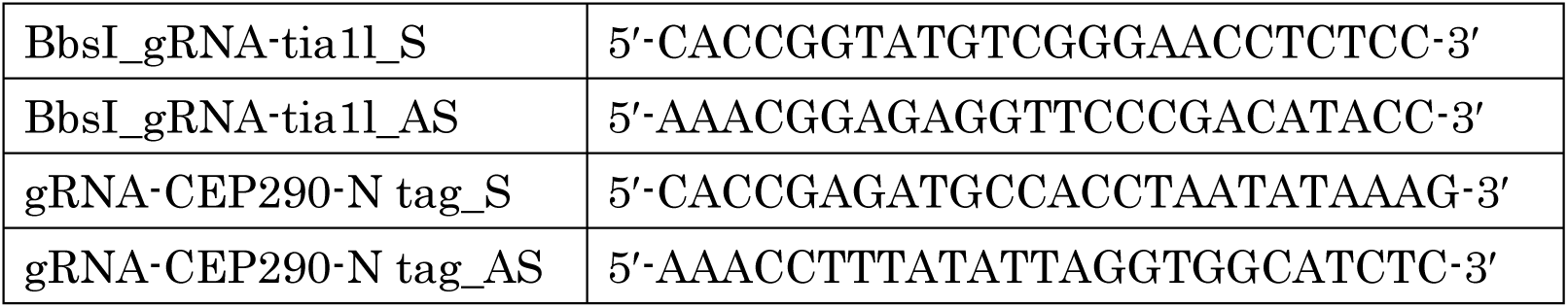

We established a PA-tagged CEP290 cell with Non homologous end joining (NHEJ)-based method, as previously described with modifications [17] (see Supplemental Fig. 1). Briefly, using the synthesized oligonucleotides, the sgRNA template was cloned into pX459 and designated as pX459_sgRNA-CEP290-N_sgRNA-tia1l. The NHEJ knock-in donor plasmid was constructed and designated as pDonor-N-Hygro^R^-T2A-PA tag. These two plasmid constructs were transfected into mIMCD3 cells, and drug-resistant clones were isolated under 2 μg/mL of puromycin and 25 μg/mL of hygromycin D. The genotype of the isolated clone was confirmed by sequencing genomic PCR products. The primers used were as follows:

Oligonucleotides for CRISPR/Cas9 mediated gene targeting

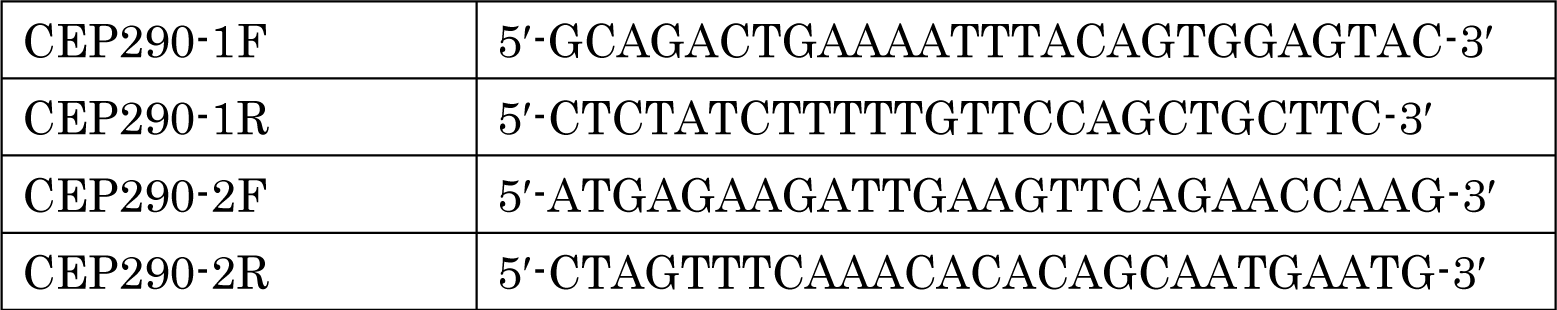

PCR/Sequencing primers for genotyping were as follows:

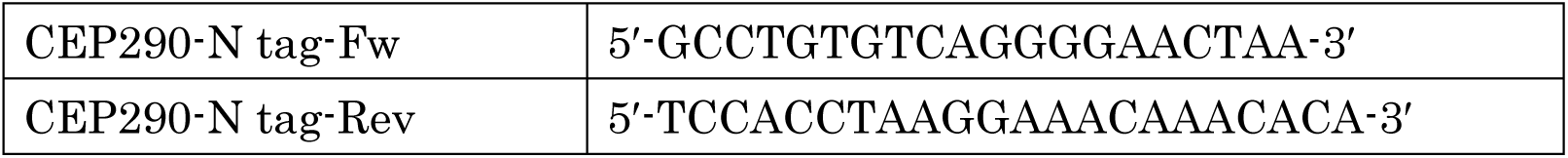

### 2.2 Antibodies and immunostaining

Antibodies used in this study were as follows:

2.2 Antibodies and immunostaining Antibodies used in this study were as follows:
| Antibody | Cat. No. | Industry | dilution factors |
| --- | --- | --- | --- |
| Anti-CEP290 rabbit polyclonal antibody | NB100-86991 | Novus Biologicals, Centennial, CO, USA | x500 (WB), x300 (ICC) |
| Anti-p150<sup>Glued</sup> mouse monoclonal antibody | P41920 | BD Bioscience, Franklin Lakes, NJ, USA | x1000 (WB), x500 (ICC) |
| Anti-α-tubulin mouse monoclonal antibody, clone DM1A | 05-829 | Merck, Darmstadt, German | x1000 (ICC) |
| Anti-α-tubulin rabbit polyclonal antibody | PM054 | MBL, Tokyo, Japan | x500 (ICC) |
| Anti-γ-tubulin mouse monoclonal antibody, clone GTU88 | T5326-200U | Merck | x300 (ICC) |
| Anti-paxillin mouse monoclonal antibody, clone 5H11 | 05-417 | Merck | x500 (ICC) |
| Anti-PA rat monoclonal antibody | 012-25863 | Fujifilm, Tokyo, Japan | x1000 (WB), x500 (ICC) |
| Anti-APC rabbit polyclonal antibody | 2504S | Cell Signaling Technology, Danvers, MA, USA | x1000 (WB), x500 (ICC) |
| Goat anti-rabbit IgG antibody-Alexa Fluo 488 | A-11008 | Thermo Fisher Scientific | x5000 (ICC) |
| Goat anti-mouse IgG antibody-Alexa Fluo 488 | A-11001 | Thermo Fisher Scientific | x5000 (ICC) |
| Goat anti-mouse IgG antibody-Alexa Fluo 555 | A32727 | Thermo Fisher Scientific | x5000 (ICC) |
| Goat anti-mouse IgG antibody-Alexa Fluo 633 | A-21052 | Thermo Fisher Scientific | x5000 (ICC) |
| Goat anti-rat IgG antibody-Alexa Fluo 555 | A48263 | Thermo Fisher Scientific | x5000 (ICC) |

WB: Western blotting, ICC: Immunocytochemistry

Alexa Fluo 633 Phalloidin was used for F-actin staining (A22284; Thermo Fisher Scientific). Nuclei were stained with 4′,6-Diamidino-2-phenylindole, dihydrochloride (DAPI) (340-07971: DOJINDO, Kumamoto, Japan).

For immunostaining, cells were fixed with 4% paraformaldehyde in phosphate buffered saline (PBS) without divalent cations for 30 min at 25 °C or in methanol for 5 min at -20 °C. After removing the fixative agent by washing with PBS, the specimens were immersed in a blocking solution (PBS containing 5% normal goat serum, 0.03% TritonX-100 and 0.1 % sodium azide) at 25 °C for 60 min. Subsequently, a primary antibody was applied to the specimens at an indicated dilution in blocking solution. After incubation for 60 min at 25 °C, the redundant antibody was removed by washing with PBS-T (PBS containing 0.03% Triton X-100), a secondary antibody, and DAPI appropriately diluted in the blocking solution and was applied to the specimens. After 30 min of incubation at 25 °C, the redundant antibody and DAPI were removed by washing with PBS-T. The specimens were mounted using FluorSave reagent (345789; Merck). Images were acquired using an LSM900-Airyscan confocal laser scanning microscope (ZEISS, Oberkochen, German). Line profiles were processed using Zen software (ZEISS).

### 2.3 Microtubule re-growth assay

The microtubule re-growth assay was performed as previously described [18]. Briefly, wild-type CEP290-KO1 and CEP290-KO2 cells were cultured on glass coverslips overnight (O/N) and then incubated on ice for 30 min to depolymerize the microtubule fibers. Next, cold cultural medium was replaced with 37 °C-prewarmed medium. The cells were immediately transferred to a CO_2_ incubator at 37 °C and incubated for 3 min. The cells were subsequently fixed with cold methanol (−20 °C), followed by staining with anti-α-tubulin antibody (DM1A: #05-829, Merck). The specimens were observed under an IX83 fluorescence microscope (Evident, Tokyo, Japan). The acquired images were processed using CellSence Dimensions (Evident) and ImageJ/Fiji software.

### 2.4 Cell migration assay

We seeded 5 × 10^4^ cells of wild-type or CEP290-KO into a Culture-Insert 2 well (ib80209, ibidi GmbH, Bayern, German) set onto a well of the 24 well plate (3820-024, Iwaki, Tokyo, Japan) to make a cell free gap of 500 μm. Culture inserts were removed 24 hours after seeding, and cell migration was observed using an In Cell Analyzer 2200 (Cytiva) every 6 h for 24 h.

### 2.5 Cell attachment assay and nocodazole treatment

For specimen preparation, 1 × 10^5^ cells were seeded in a 6-well plate (140675, Thermo Fisher Scientific) in which glass coverslips coated with collagen IV (Nitta Gelatin, Inc., Osaka, Japan) were used. For the cell attachment assay, the culture medium was replaced with PBS and weakly adherent cells were removed by washing with PBS after 4 h of incubation. The cells remaining on the glass coverslips were fixed with 4% paraformaldehyde (PFA) in PBS. Immunostaining was performed as previously described. Nocodazole treatment included adding nocodazole to the growth medium at a final concentration of 0.1 ng/mL 4 h after seeding onto collagen IV-coated coverslips. The cells were incubated at 37 °C for 3 min, weakly adherent cells were removed by washing with PBS, and the remaining cells were subsequently fixed with 4% PFA. Fixed cells were subjected to immunostaining. Images were acquired using APEXVIEW APX100 (Evident). Cell and paxillin areas were measured using Fiji software [19]. Data were statistically analyzed in triplicates.

### 2.6 3D live imaging of EB3-EGFP

Full-length cDNA of human EB3 was amplified by RT-PCR from total RNA harvested from hTERT-RPE1 (retinal pigment epithelial) cells, and the PCR products were cloned into the *Eco*RI/*Sal*I multicloning sites of the pEGFP-N3 vector (Takara), designated as pEGFP-hEB3. pEGFP-hEB3 was transfected with the Trans-IT LT1 reagent (Takara) into wild-type or CEP290-KO cells seeded in a glass-bottomed dish (Matsunami Glass Ind.). Live imaging data were acquired by Zeiss LSM900-Airyscan confocal fluorescent microscope. The velocity of the EB3-EGFP positive foci was measured using Imaris software (Oxford Instruments). hTERT-RPE1 cell line was obtained from ATCC (CRL-4000).

### 2.7 Proximal ligation assay

The proximal ligation assay (PLA) was performed using the Duolink In Situ Orange Starter Kit Mouse/Rabbit (DUO92103; Sigma-Aldrich), according to the manufacturer’s protocol. Briefly, the cell specimens fixed by 4% PFA were treated with Duolink blocking solution for 60 min at 37 °C. Next, an anti-APC antibody (rabbit), appropriately diluted in Can Get Signal A solution (TOYOBO, Osaka, Japan), was applied to the specimens. The specimens were washed twice with wash buffer A after incubation for 60 min at 25 °C. Anti-PA tag-PLUS PLA probe and anti-rabbit IgG-MINUS PLA probes (DUO92005: Sigma-Aldrich) appropriately diluted in the Duolink antibody diluent were applied onto the specimens that were incubated for 60 min at 37 °C. Anti-PA-tagged antibody conjugated with a PLUS oligonucleotide was prepared using the Duolink In Situ Probemaker PLUS (DUO92009; Sigma-Aldrich). The specimens were washed twice with wash buffer A and were subsequently incubated with Duolink ligation solution for 30 min at 37 °C. After the ligation reaction, the specimens were washed twice with wash buffer A and immersed in an amplification buffer containing polymerase.

Amplification reaction was performed by incubation in a pre-heated humidity chamber for 100 min at 37 °C. The specimens were then washed with wash buffer B and mounted for observation. Data were acquired using an IX83 immunofluorescence microscope, and the obtained images were processed using CellSense Dimension.

## 3. Results

### 3.1 *Cep290* is essential for MTOC function of the centrosome

To elucidate the molecular functions of CEP290 in non-ciliated cells, we generated *Cep290* knockout cell line using the CRISPR-Cas9 system in mIMCD3 cells. To avoid misidentifying the off-target effect as a specific phenotype, we generated two independent *Cep290* null alleles, CEP290-KO1 and CEP290-KO2, using differently designed sgRNAs (Fig. 1A). Protein production of CEP290 was completely missing in both CEP290-KO1 and -KO2 cells (Fig. 1B). In non-ciliated cells, CEP290 localizes to the centrosome [6,13,14]. Because the centrosome is a major MTOC in mammalian cells, we examined whether *Cep290* is required for MTOC function using a microtubule regrowth assay. The microtubule fibers were well recovered throughout the cytoplasm of the wild-type cells 3 min after the cells were returned to the regrowth condition. Contrastingly, the microtubule fibers remained disrupted in all CEP290-KO cell lines, and tubulin protein was concentrated around the centrosome (Fig. 1C). Furthermore, we investigated the microtubule dynamics in *Cep290* mutants. To measure the velocity of the microtubule plus end, a member of the microtubule plus end-tracking proteins (+TIPs) tagged with GFP, EB3-EGFP, was transiently expressed [20]. The moving velocity of EB3-EGFP represents the dynamics of microtubule growth. We subsequently found that the velocity of EB3-EGFP was significantly slower in the CEP290-KO cells than that in the wild-type cells (Fig. 1D and supplemental movie 1-3). Thus, CEP290 is essential for MTOC function and the proper growth of microtubules.

**Fig.1.**
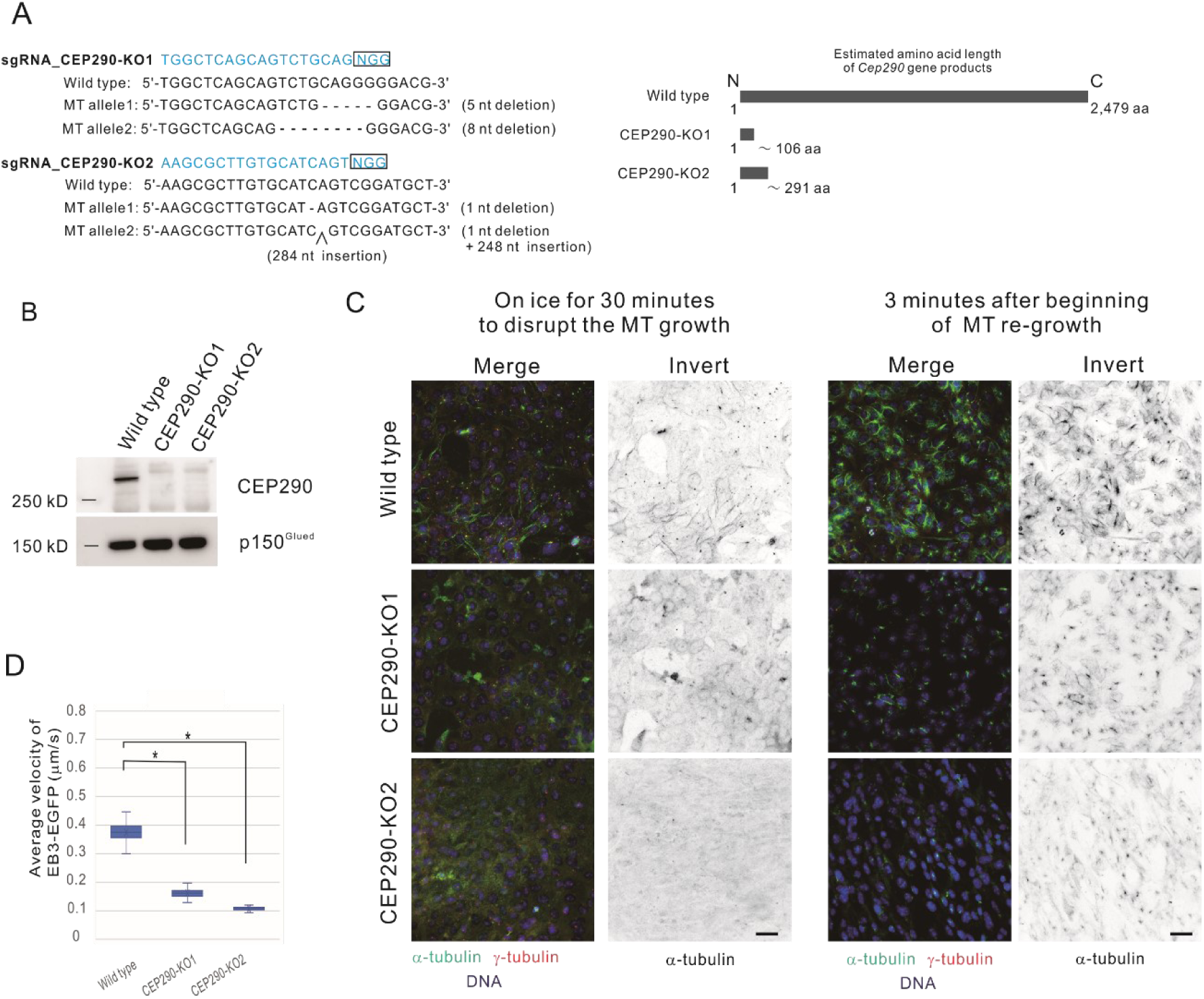
CEP290 knockout impairs microtubule organizing center (MTOC) function of the centrosome. A. CEP290-KO1 and CEP290-KO2 allele. The boxed sequences in the left panel represent protospacer adjacent motif (PAM). The right panel shows expected amino acid length of the products from the mutant alleles. B. The production of CEP290 in wild-type and CEP290-KO cells was analyzed by western blotting. C. Microtubule regrowth assay. Incubation of the wild-type and CEP290-KO cells on ice induced depolymerization of almost all microtubule fibers, and tubulins were seen only at the centrosome in the wild-type and CEP290-KO cells. After restoring the temperature to 37 °C, microtubule fiber was regrown from the centrosome in wild-type cells, whereas significant regrowth of microtubule fibers was not observed in CEP290-KO cells. Green: α-tubulin, Red: γ-tubulin, and Blue: DNA. A monochrome picture represents inverted image of α-tubulin staining. Scale bar indicates 40 μm. D. The box-plot represents EB1-EGFP velocity in the wild-type or CEP290-KO cells. An asterisk indicates statistical significance (p < 0.01).

### 3.2 Malfunction of CEP290 affects directed cell migration

The proper regulation of microtubule dynamics is critical for directed cell migration. MTOC malfunction results in abnormal microtubule dynamics. We subsequently investigated whether directed cell migration of *Cep290* knockout cells was impaired using a cell migration assay. We monitored how a gap of 500 μm width on a cell monolayer was being filled by cell migration. Compared to the wild-type cells, the CEP290-KO cells took longer to fill this gap (Fig. 2). Thus, as expected, CEP290 contributed to directed cell migration through microtubule dynamics, likely by regulating MTOC function.

**Fig.2.**
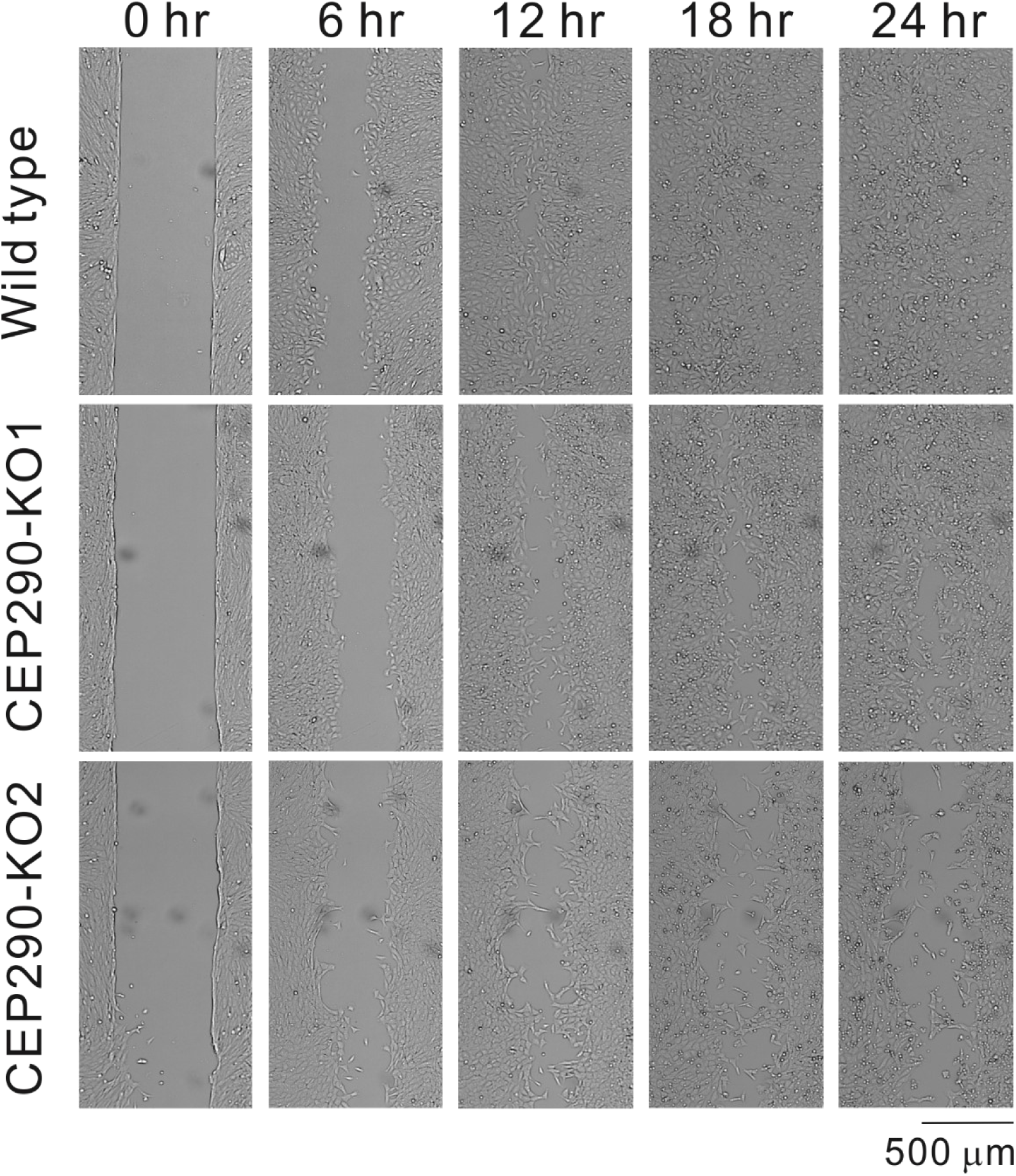
CEP290 knockout impairs directed cell migration. Data from the cell migration assays are shown. The initial gap width is 500 μm. The cells were monitored for up to 24 h. Notably, filling of the gaps in mutations takes time. Scale bar indicates 500 μm.

### 3.3 Cell adhesion depends on functional CEP290

Microtubules have been considered to play an important role in cell migration. However, the underlying molecular mechanism remains unclear. Generally, microtubules are involved in cell migration by modulating actomyosin dynamics, transporting the required membrane components to the leading edge, and/or supplying molecules related to cell adhesion to adhesion plaques [21]. In our experiment, we observed that the mutant cells detached easily from the culture dish surface. This evidence suggests that *Cep290* regulates cell adhesion to the extracellular matrix (ECM). To test this hypothesis, we assayed the adhesion of mutant cells to type IV collagen, a component of the basement membrane, in comparison with the wild-type cells. We seeded the determined number of cells on the glass surface coated with type IV collagen, incubated them for 4 h, washed and removed the cells that did not sufficiently adhere, and observed how efficiently the disseminated cells were attached to the surface (Fig. 3 A and B). Consequently, the ability of the mutant to adhere to the collagen IV-coated surface was lower than that of the wild-type cells. Additionally, the mutant cells were smaller and shrunken than the wild-type cells (Fig. 3A and C). To further explore this hypothesis, we examined the status of the cell adhesion plaques. The distribution of the main component of focal adhesion, paxillin, was clearly reduced in the mutants, particularly in the leading-edge region (Fig. 3D and E). Thus, CEP290 may control focal adhesions by regulating MTOC and/or microtubule dynamics.

**Fig.3.**
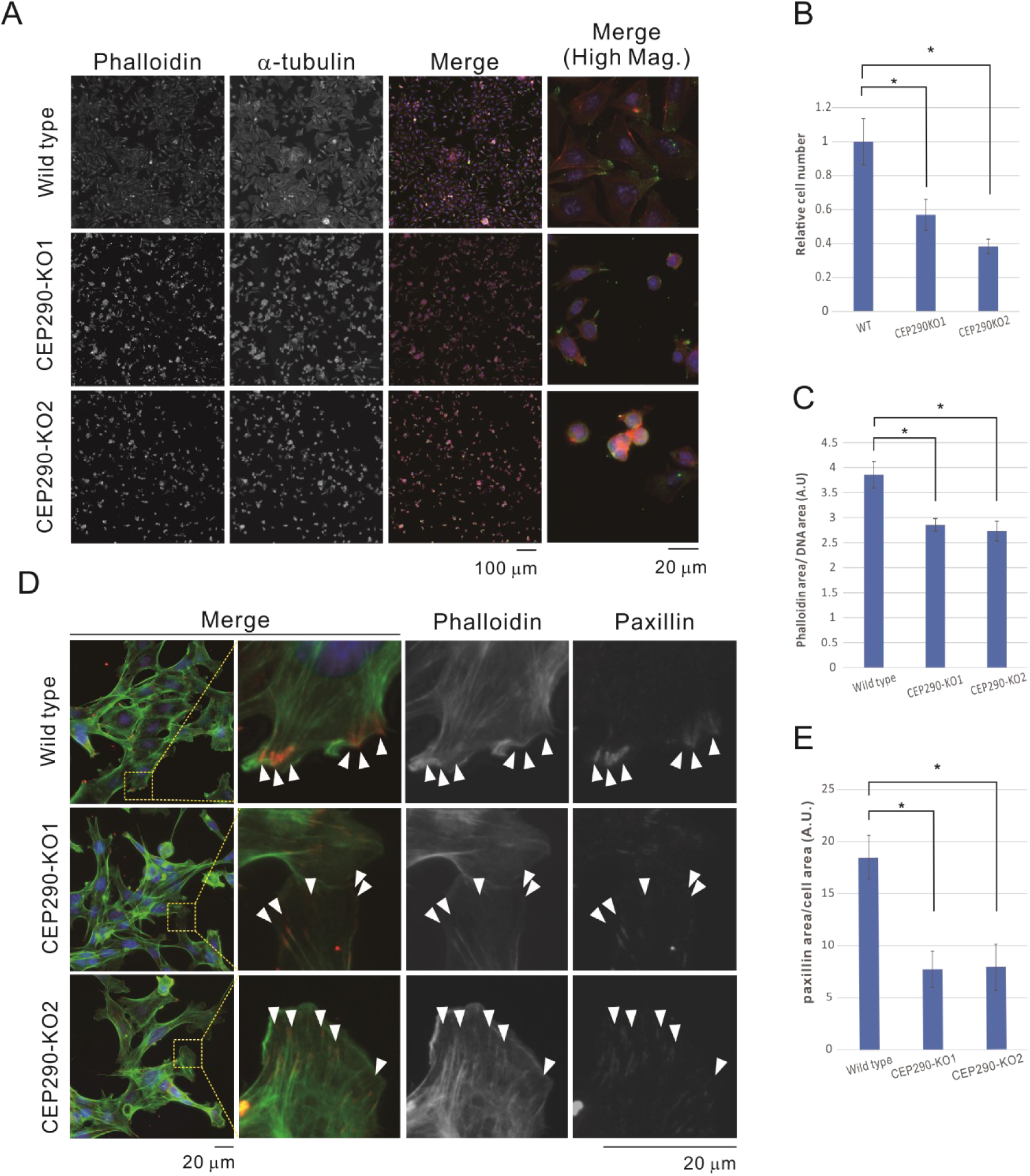
Reduced adhesive capacity in CEP290 knockout cells. A. The knockout cells were shrunken and less adhesive. The determined cell number were seeded on the glass coverslips coated with collagen IV. The cells that did not sufficiently adhere on the coverslips were removed 4 h after seeding, and the cell number and morphology were observed. Magnified images of a typical area of each cell are displayed on the far right. Red: F-actin stained with Phalloidin, Green: α-Tubulin, and Blue: nucleus). A scale bar indicates 100 μm and 20 μm in lower and higher magnifications, respectively. B. The counted cell number in the same experiment as in A indicates the mutant as less adhesive. An asterisk indicates statistical significance (p < 0.01). Wild-type number is set to one in this graph. C. CEP290 mutant cells became shrunken. The graph shows the area occupied by cells normalized by the area occupied by the nucleus. An asterisk indicates statistical significance (p < 0.01). D. Focal adhesion formation of the mutant was reduced in the mutant cells. Green: F-actin, Red: paxillin, and Blue: nucleus. Scale bar indicates 20 μm. The boxed areas are magnified on the right. Arrow heads indicate focal adhesions. E. The area occupied by paxillin normalized by the cell area in D was measured. An asterisk indicates statistical significance (p < 0.01). Note that the focal adhesion was significantly reduced in the mutants.

### 3.4 APC-CEP290 complex transports paxillin to the leading-edge via a microtubule network

To further explore the relationship between CEP290 and paxillin, we tested whether CEP290 binds directly to paxillin. Unfortunately, we could not find any commercially available anti-CEP290 antibody for immunostaining and immunoprecipitation (IP) of specimens from mIMCD3 cells. Alternatively, we genetically modified the *Cep290* gene in mIMCD3 cells using the CRISPR-Cas9 system, as the N-terminus of CEP290 possesses a PA tag (PA-CEP290). We validated PA-CEP290 production by western blotting (Fig. 4A) and its subcellular localization by immunostaining (Fig. 4B). As expected, we found that PA-CEP290 was endogenously produced and localized to the centrosome [6,14]. Furthermore, we did not observe any abnormality in the PA-CEP290 mIMCD3 cell line in morphology, growth, migration, or adhesive capacity (data not shown). Thus, we conclude that PA-CEP290 functions physiologically similar to endogenous molecules. Using this cell line, we performed an IP assay with anti-PA antibody or anti-paxillin antibody, but we did not observe any co-immunoprecipitated PA-CEP290 with paxillin by western blotting (data not shown). This strongly suggested that CEP290 does not directly bind to paxillin.

**Fig.4.**
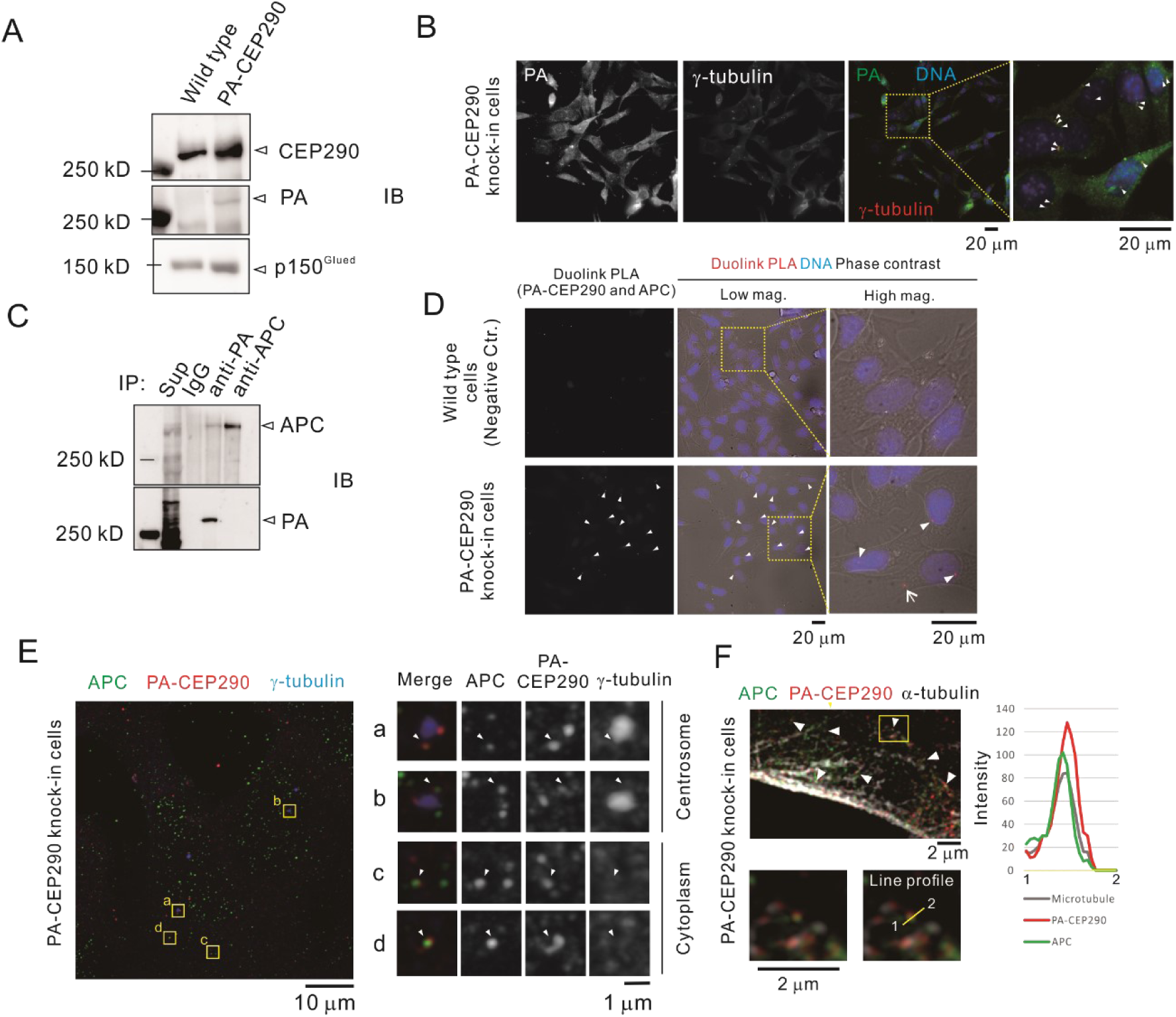
CEP290 forms a complex with adenomatous polyposis coli (APC) on the microtubule network. A. PA-CEP290 cells in which N-terminal of CEP290 is tagged with PA tag were generated via CRISPR/Cas9. Western blot shows PA-CEP290 identified by anti-PA antibody. p150^Glued^ was used as a loading control. B. The subcellular distribution of PA-CEP290 was confirmed by immunostaining. Green: PA-CEP290, Red: γ-Tubulin, and Blue: nucleus). Scale bars indicate 20 μm. Note that PA-CEP290 localization at the centrosome represented by γ-tubulin is as expected (white arrow heads). C. Interaction between CEP290 and APC were analyzed by co-IP. Anti-PA antibody allows co-IP of PA-CEP290 with APC, although anti-APC antibody did not allow co-IP of APC with PA-CEP290. See main text in detail. D. Red color signal in proximal ligation assay (PLA) shows co-localization of PA-CEP290 and APC (white arrow heads). Scale bars indicate 20 μm. The boxed areas are magnified on the far right. The PLA signal indicated by arrowhead is the centrosome, and the signal that was not related to the centrosome was also observed in the cytoplasm (arrow). E. Immunofluorescence of APC (green), PA-CEP290 (red), and γ-tubulin (blue) on PA-CEP290 cells observed with super resolution fluorescence microscope. The scale bar indicates 10 μm and 1 μm in lower and higher magnifications, respectively. Region of interest (ROI) of a to d (yellow squares) are magnified on the right. ROI of a and b showed co-localization of APC and PA-CEP290 at the centrosome (γ-tubulin). Note that the ROI of c and d showed co-localization in the cytoplasm not related to the centrosome. F. Super resolution immunofluorescence of APC (green), PA-CEP290 (red), and α-tubulin (grey) on PA-CEP290 cells. A yellow boxed area in the upper panel is magnified in the lower panel. The fluorescent intensities of APC, PA-CEP290, and α-tubulin on an indicated yellow line in the lower panel were measured by line profile. Note that all peak positions of the fluorescence intensities of APC, CEP290, and α-tubulin were approximately the same. Scale bars indicate 2 μm.

Contrastingly, a previous study showed that APC, a microtubule-associated protein and regulator of cell adhesion, is vital for paxillin localization at the leading edge [22]. We hypothesized that APC and CEP290 form a complex that transports paxillin to the focal adhesion or leading edge. To examine this, we first assayed the interaction between CEP290 and APC by IP. We found that APC co-immunoprecipitated with PA-CEP290 when an anti-PA antibody was used for IP (Fig. 4C), suggesting that CEP290 forms a complex with APC. However, when the anti-APC antibody was used for IP, the PA-CEP290 band was not detected by western blotting. This result may be a consequence of the existing ratio of APC to CEP290 molecules, that is, the APC number is much higher than that of CEP290, and an exceedingly small portion of APC molecules in a cell contribute to the APC-CEP290 complex.

To further confirm the complexity of APC-CEP290, we investigated whether APC and CEP290 molecules closely co-localize in cells. The proximal ligation assay, which can detect and visualize localization only when the molecules of two different analysis targets are in close proximity, showed that CEP290 and APC interact at the centrosome and in the cytoplasm (Fig. 4D). Importantly, using super-resolution microscope we found co-localization of CEP290 and APC sparsely throughout the cytoplasm in addition to the co-localization of PA-CEP290 and APC in the proximity of centrosome represented as the γ-tubulin foci, although many APC molecules unrelated to CEP290 were found (Fig. 4E). This evidence indicate that APC-CEP290 truly forms a complex, but only a small number of APC molecules interact with CEP290. Notably, we observed that the fluorescent signals of APC, PA-CEP290, and α-tubulin overlapped, suggesting that APC-CEP290 complexes are on the microtubule fibers (Fig. 4E and F). Line profile analysis indicated that the APC, CEP290, and α-tubulin signals were spatially located in approximately the same place (Fig. 4F). This finding suggests that APC-CEP290 complexes exist in the centrosome and on microtubule fibers. This complex is transported via a microtubule network. Given that APC binds to paxillin and plays an important role in its distribution [22], CEP290 seems to regulate paxillin transport via APC.

### 3.5 Focal adhesion formation does not depend on dynamic instability of microtubules but trafficking of APC-CEP290 complexes

Nocodazole, which binds to β-tubulin and disrupts microtubule assembly/disassembly dynamics, enhances stabilization and area expansion of focal adhesion by suppressing microtubule dependent turnover of focal adhesion-related molecules [23]. Therefore, we investigated whether nocodazole treatment could rescue the phenotype of reduced adhesion capacity and focal adhesion formation in the *Cep290* knockout (Fig. 5A-C). First, we observed that the intensity of the intracellular paxillin signal increased in the wild-type and mutant cells after nocodazole treatment (Fig. 5C). This result, which was expected based on a previous report, supports the notion that nocodazole works accurately. However, the CEP290-KO cell phenotypes, such as the shrunken cell shape, reduced size and focal adhesion number, and weak adherence ability to the culture dish, were not rescued (Fig. 5A and B). Thus, the paxillin accumulation in the adhesion plaques is insufficient for cell adhesion if the focal adhesions are reduced in size and number.

**Fig.5.**
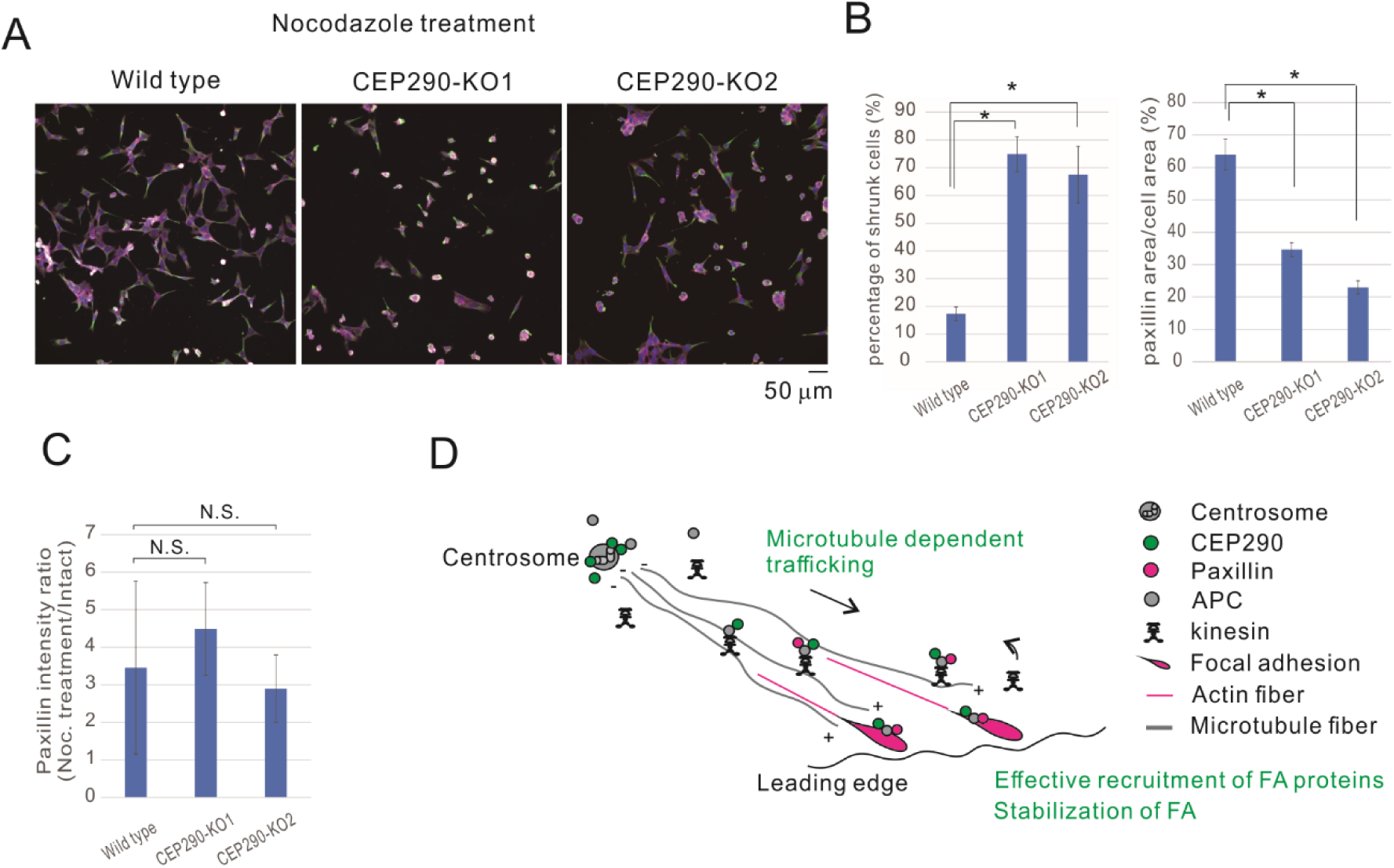
CEP290 transports paxillin to focal adhesion/leading edge via a supporting microtubule growth and complex with adenomatous polyposis coli (APC) A. The same experiment as in Fig. 3A was performed with nocodazole treatment. Note that this did not rescue the less adhesive feature of the *Cep290* mutant. Red: F-actin, Green: microtubule, and Blue: nucleus. Scale bar indicates 50 μm. B. Nocodazole treatment did not rescue focal adhesion formation. The shrunken cell percentage was not reduced by nocodazole in the mutants (left panel). Importantly, the area of focal adhesion marked by paxillin was still significantly lower in the mutants than in the wild-type (right panel). An asterisk indicates statistical significance (p < 0.01). C. The graph shows the ratio of paxillin intensity with nocodazole normalized by that without nocodazole. The paxillin signal is enhanced by nocodazole treatment in all cell types. This indicates that nocodazole inhibited the turnover of paxillin as expected. N.S.: no statistical significance. D. A schematic model of CEP290 for regulating focal adhesion (refer to the main text).

## 4. Discussion

Primary cilia are important for transmitting extracellular signals into a cell. Ciliary protein dysfunction may cause ciliopathies. Generally, ciliopathy phenotypes are diverse. Although some phenotypes are common to ciliopathies, others are ciliopathy-specific.

In our study, we hypothesized that some phenotypes of ciliopathies depend directly on primary cilia, whereas others depend indirectly. CEP290 is an important ciliary protein involved in regulating ciliary signaling. Mutations in *Cep290*, the gene causing ciliopathy, result in various phenotypes, possibly due to the complex interplay between primary ciliary dysfunction and other dysfunctions. We analyzed CEP290 function in non-ciliated cells not related to primary cilia. In summary, CEP290 is required for regulating MTOC, directed cell migration, and cell adhesion to the ECM (Fig. 5D). Focal adhesion formation dependent on CEP290 seems vital for directed cell migration and ECM adhesion. CEP290 forms a complex with APC and likely transports the paxillin molecule toward the focal adhesion/leading edge by binding to the APCs. CEP290 also governs microtubule growth, possibly contributing to the transport of focal adhesion molecules. These findings strongly suggest that cell migration, microtubule growth, and/or cell adhesion are responsible for some CEP290-related ciliopathy phenotypes.

Given that APC is involved in WNT signaling, CEP290 may play a role in WNT signaling. In non-ciliated cells, canonical WNT signaling stabilizes cytoplasmic β-catenin, resulting in increased nuclear β-catenin levels. Nuclear β-catenin initiates the transcription of TCF/LEF1-responsive genes. In this context, if WNT signaling is inactive, the Axin-GSK3β-APC destruction complex interacts with β-catenin and promotes its degradation, thereby suppressing the WNT signaling pathway. We did not clarify whether CEP290 is involved in WNT signaling via APC, but we cannot exclude its possibility. However, further studies are required to address this issue.

Unexpectedly, our study showed that the effective trafficking of paxillin to focal adhesion requires a molecule that seems to play an important role at the centrosome and cilium. However, previous studies have revealed that (1) paxillin [24] and (2) APC localize at the centrosome, plays a significant role in microtubule growth, and directly binds to paxillin and focal adhesion kinase [22,25–28]. Therefore, it is reasonable to conclude that the centrosome molecule CEP290 is directly involved in paxillin regulation. Interestingly, nocodazole treatment to suppress paxillin turnover did not restore the impaired cell adhesion capacity of CEP290 mutants (Fig. 5A-C). Even though the paxillin molecules that are highly abundant are maintained in focal adhesions, proper cell adhesion to the ECM cannot be maintained if focal adhesion formation is reduced in size and number, similar to *Cep290* mutants.

Issues related to primary cilia have been extensively studied in terms of ciliopathy pathogenesis, although our understanding of the molecular pathogenesis of ciliopathies is still incomplete. However, our study provides a novel platform to deepen our knowledge of the function of non-primary cilia in ciliopathy-related molecules. Further analyses are needed to clarify the roles of primary ciliary proteins in primary cilia and in structures other than primary cilia to validate the molecular pathogenesis of ciliopathies.

## Supporting information

Supplemental informations

## Acknowledgements

We thank the Kyoto University Live Imaging Center (KULIC) for the imaging data acquisition. We would like to thank Editage (www.editage.com) for English language editing.

## ^1^Abbreviations

APC: adenomatous polyposis coli
CEP290: centrosomal protein 290
MTOC: microtubule organizing center
mIMCD3: Murine inner medullary collecting duct 3
NHEJ: Non homologous end joining
+TIPs: plus end-tracking proteins
PLA: Proximal ligation assay

## Figure Legends

