## Supplemental informations for "Ciliary protein CEP290 regulates focal adhesion via microtubule system in non-ciliated cells^1^"

Supplementary Material

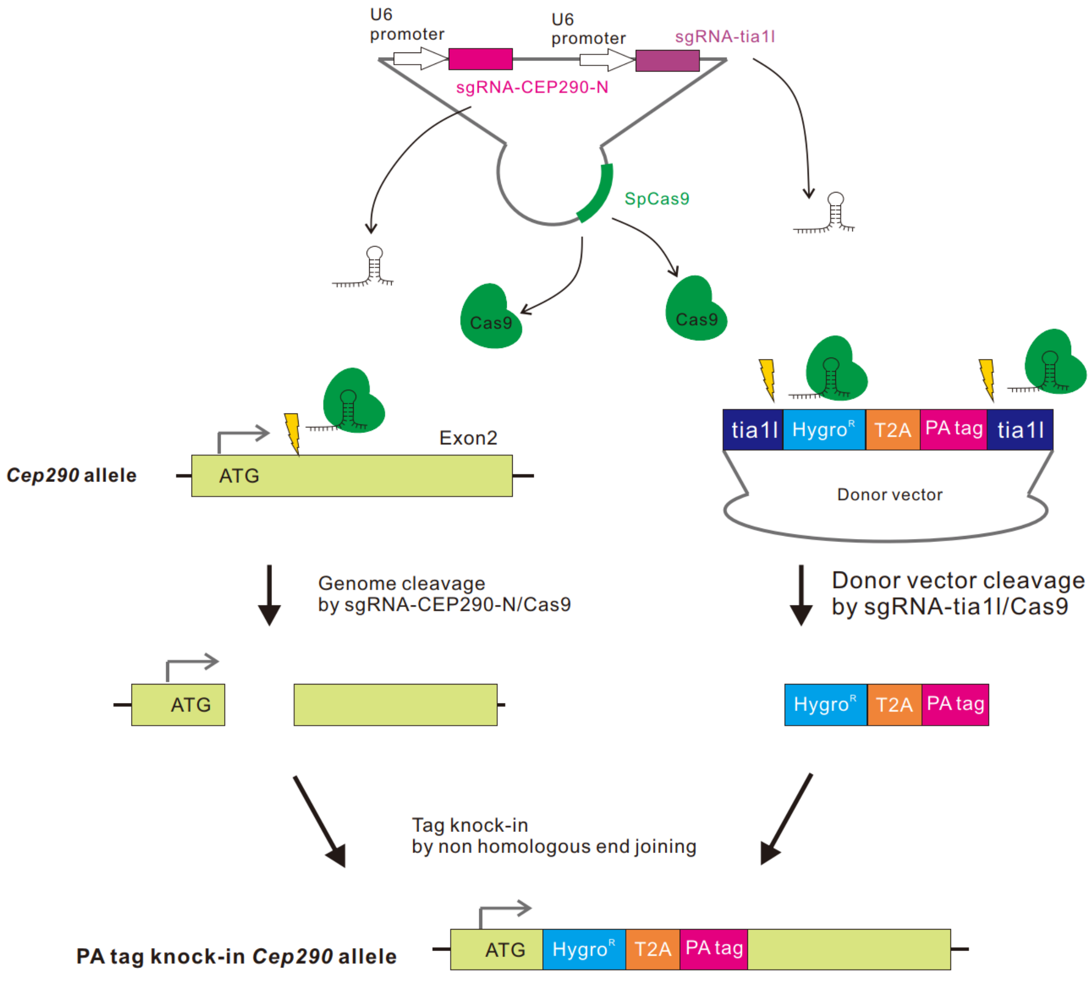


Supplemental Fig.1

Supplemental Fig.1. The strategy of tagging CEP290 with PA tag.

We designed sgRNA-CEP290-N and sgRNA-tia1l to create a HygroR-T2A-PA tag cassette knocked-in in-frame into the *Cep290* gene, following the first methionine codon (ATG) of the ORF by non-homologous end joining (NHEJ). The *Cep290* allele was excised from the Cas9/sgRNA-CEP290-N complex. The donor vector containing the Hygro$^{R}$-T2A-PA tag cassette flanked by two tia1l sequences was cut using the Cas9/sgRNA-tia1l complex.

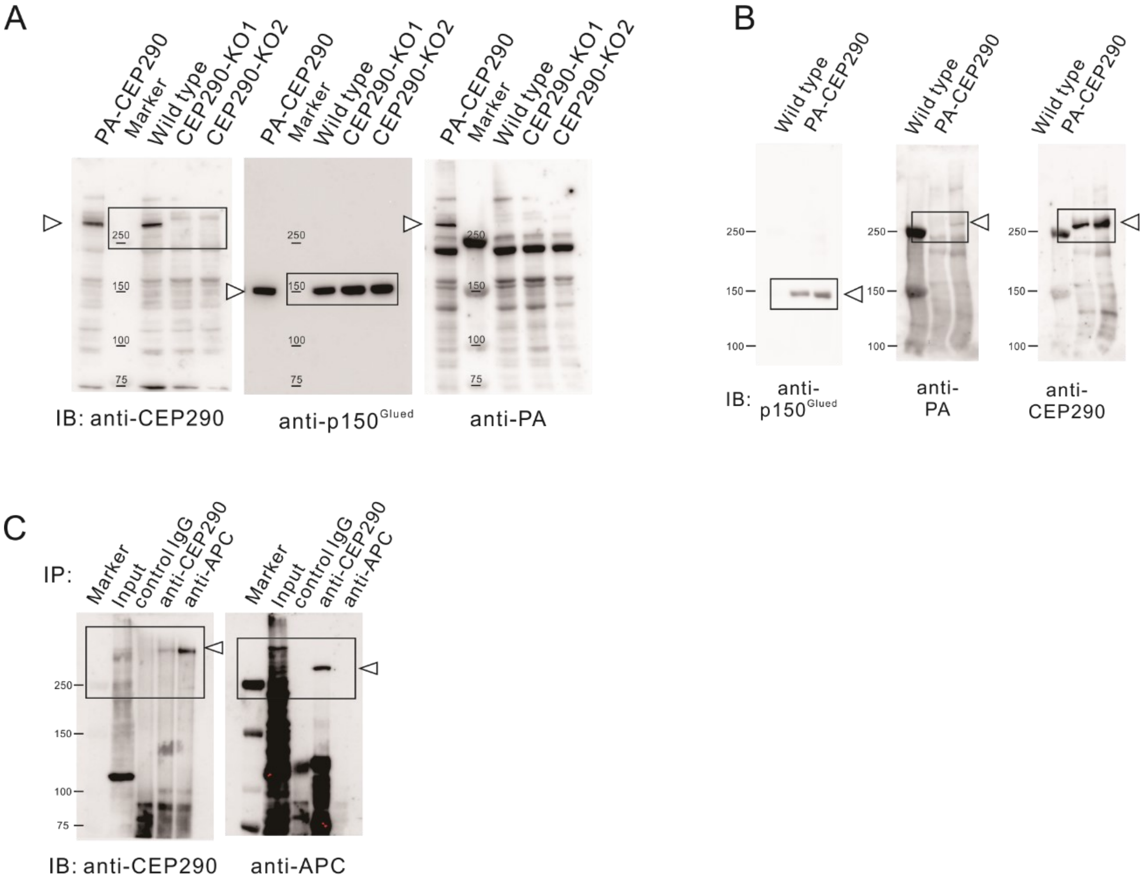


Supplemental Fig.2

Supplemental Fig.2. Whole gel images of western blot used in this study.

The boxed areas in the gel images of (A), (B), and (C) are shown in Figs. 1B, 4A, and 4C, respectively.

Supplemental movies

Supplemental movie1. Live imaging data of EB3-EGFP in the wild-type cell.

As shown in Fig. 1D, the 3D live imaging data of EB3-EGFP in wild-type cells were orthogonally projected. Scale bar indicates 5 μm.

Supplemental movie2. Live imaging data of EB3-EGFP in CEP290-KO1 cell.

As shown in Fig. 1D, 3D live imaging data of EB3-EGFP in CEP290-KO1 cells were orthogonally projected. The scale bar indicates 5 μm.

Supplemental movie3. Live imaging data of EB3-EGFP in CEP290-KO2 cell.

As shown in Fig. 1D, 3D live imaging data of EB3-EGFP in CEP290-KO2 cells were orthogonally projected. Scale bar indicates 5 μm.
